# Shed skin as a source of DNA for genotyping-by-sequencing (GBS) in reptiles

**DOI:** 10.1101/658989

**Authors:** Thomas D Brekke, Liam Shier, Matthew J Hegarty, John F Mulley

## Abstract

Over a fifth of reptile species are classified as ‘Threatened’ and conservation efforts, especially those aimed at recovery of isolated or fragmented populations, will require genetic and genomic data and resources. Shed skins of snakes and other reptiles contain DNA, are a safe and ethical way of non-invasively sampling large numbers of individuals, and provide a simple mechanism by which to involve the public in scientific research. Here we test whether the DNA in dried shed skin is suitable for reduced representation sequencing approaches, specifically genotyping-by-sequencing (GBS). Shed skin-derived libraries resulted in fewer sequenced reads than those from snap-frozen muscle samples, and contained slightly fewer variants (70,685 SNPs versus 97,724), but this issue can easily be rectified with deeper sequencing of shed skin-derived libraries. Skin-derived libraries also have a very slight (but significantly different) profile of transitions and transversions, most likely as a result of DNA damage, but the impact of this is minimal given the large number of single nucleotide polymorphisms (SNPs) involved. SNP density tends to scale with chromosome length, and microchromosomes have a significantly higher SNP density than macrochromosomes, most likely because of their higher GC content. Overall, shed skin provides DNA of sufficient quality and quantity for the identification of large number of SNPs, but requires greater sequencing depth, and consideration of the GC richness of microchromosomes when selecting restriction enzymes.

## Introduction

The first extinction-risk assessment of reptiles was published in the spring of 2022, and showed that just over a fifth (21.1%) of reptile species are threatened, making them the second most vulnerable group after amphibians (Cox et al., 2022; ‘The IUCN Red List of Threatened Species’). Conserving these species will require interventions at a variety of levels, including not only habitat preservation and restoration, but also the development of data and resources for the estimation of genetic factors associated with extinction risk and recovery potential. Genetic tools are important for the identification and delineation of species (Paul D. N. Hebert et al., 2003; Paul D N Hebert et al., 2003); assessment of gene-flow and sex-biased dispersal; estimation of effective population size; and for determining historical changes such as range expansions and contractions (DiLeo and Wagner, 2016; Shaffer et al., 2015). Such studies have historically relied on small numbers of genetic markers such as simple sequence repeats (SSRs, microsatellites) (e.g. (McCracken et al., 1999; Scott et al., 2001; Blouin-Demers and Gibbs, 2003; Bond et al., 2005), although next-generation sequencing approaches have facilitated the identification of much larger numbers of SSRs (Castoe et al., 2012). More recently, reduced representation sequencing approaches such as restriction site-associated DNA sequencing (RADseq (Baird et al., 2008; Davey and Blaxter, 2010; Hohenlohe et al., 2010; Peterson et al., 2012)) and genotyping-by-sequencing (GBS (Elshire et al., 2011; Narum et al., 2013)) have enabled the identification and characterisation of huge numbers of single nucleotide polymorphism (SNP) markers. SSRs are more variable compared to typically diallelic SNPs, but SNPs are more abundant than SSRs, and are distributed more evenly across the genome, including in coding regions. SNPs can therefore provide not only broader genome coverage, but also greater statistical power (Morin et al., 2004; Zimmerman et al., 2020), and can differentiate between even very closely related individuals (Kleinman-Ruiz et al., 2017; Roques et al., 2019), something that can be particularly important in small, isolated, inbred populations. Conservation genomics studies require sources of DNA, and in the case of reptiles this can include a diverse set of tissues and sampling methodologies, including both invasive and non-invasive methods. Invasive sampling might include blood samples from a vein or via cardiac puncture (Brown, 2010; Eatwell et al., 2014), or collection of a small amount of tissue such as a toe or tail tip, or a scale clipping (Beebee, 2008; Maigret, 2019). These approaches have the disadvantages of requiring not only capture and handling/restraint of the animal, but also carry with them greater ethical implications and are likely to require special licensing. Non-invasive techniques are therefore to be preferred, and these can include fecal sampling, the use of road kill and/or museum specimens, cloacal or buccal swabs, or shed skins (Beebee, 2008; Miller, 2006; Jones et al., 2008; Lanci et al., 2012; Pearson et al., 2015). While non-invasive approaches are increasingly favoured, they are not without their own difficulties. Chemical preservation of museum specimens can degrade DNA, and procurement of adequate samples from roadkill specimens (while often high quality if found soon after death) is sporadic and unpredictable. Fecal samples inevitably contain large amounts of microorganisms, and DNA often degrades quickly unless samples are rapidly frozen (Jones et al., 2008). Cloacal and buccal swabbing (Beebee, 2008; Miller, 2006; Pidancier et al., 2003) is dependent on locating and restraining animals with research on venomous animals carrying particular risks. Shed skin samples can be collected without the need to actually locate and handle/restrain the animal itself, and without associated ethical and animal welfare issues. Such risk-free approaches lend themselves especially to citizen science projects, where members of the public can collect and ship samples, and indeed the Amphibian and Reptile Groups of the UK (ARG UK) and Amphibian and Reptile Conservation Trust (ARC) currently run a ‘Reptile Slough Genebank Project’ (https://www.arguk.org/get-involved/projects-surveys/the-reptile-slough-genebank), which asks members of the public to send them any shed skins they might find. DNA derived from shed skins of lizards and snakes has long been known to be of sufficient quality and quantity for PCR-based genotyping of a small number of genetic markers (Bricker et al., 1996; Fetzner Jr, 1999; Villarreal et al., 1996; Horreo et al., 2015; Tawichasri et al., 2017), and so here we assess the utility of shed skin for larger-scale single nucleotide polymorphism genotyping using genotyping-by-sequencing (GBS).

## Methods

### DNA extraction

Shed skins were collected from 61 corn snakes (*Pantherophis guttatus*) from our in-house colony, and from commercial breeders and hobbyist keepers (see Supplemental Table S1 for sample details). Skins were collected as soon as possible after shedding, and stored at −20°C. Those collected for us by others were placed into individual paper envelopes for shipping and were stored at −20°C upon arrival in Bangor. DNA was extracted from around 50mg samples of ventral scale skin using the DNeasy Blood and Tissue kit (Qiagen) according to the manufacturer’s protocol, with the exception of a longer (24 hour) proteinase treatment at 56°C. Small DNA fragments were removed by spin-column chromatography with Chroma-Spin-1000+TE columns (Clontech) following the manufacturer’s protocol, and samples were quantified using the Qubit dsDNA BR assay kit and Qubit Fluorometer. In some cases, it was necessary to perform multiple extractions from a single sample and pool them using ethanol precipitation to obtain the desired 100ng of DNA. We also prepared DNA from 50mg samples of snap-frozen muscle from a further 18 individuals using the same procedure.

### Genotyping-by-sequencing

Libraries were prepared as described by Elshire et al. (Elshire et al., 2011), at the Institute of Biological, Environmental and Rural Sciences (IBERS) at Aberystwyth University. Restriction digestion was carried out using the type II restriction endonuclease PstI (cut site CTGCA^G), and the resulting fragments were tagged with unique barcodes of varying lengths (Supplemental Table S1). GBS libraries were pooled to an equimolar concentration and single-end sequenced on one lane of Illumina HiSeq 2500.

### Bioinformatics

We used the stacks (v1.44) pipeline (Catchen et al., 2011, 2013) to do a genome-guided stacks assembly and call SNPs. Our pipeline started with the program ‘process_radtags’ and took as arguments the single fastq file (-f), the list of barcodes (-b), the restriction enzyme (-e pstI), as well as the flags to clean the data (-c), discard reads with low quality (-q), rescue the barcodes (-r), specify the quality encoding (-E phred33), and specify how the barcodes were situated in the reads (--inline_null). Once process_radtags had finished de-multiplexing and cleaning the data, we counted the remaining high-quality reads for each barcode. We aligned all reads to the corn snake genome (GCA_001185365.1 (Ullate-Agote et al., 2014)) with BWA (Li and Durbin, 2009) and counted the depth of coverage with ‘samtools depth’ (Li et al., 2009). We processed the genome alignments with ‘pstacks’ using a minimum stack depth of 3 (-m 3), and then built the catalog with ‘cstacks’ using all individuals and allowing for 1 mismatch against the reference (-n 1), and then ran ‘sstacks’ with all default settings. Finally, we treated all individuals as belonging to a single group and ran ‘populations’ to call variants. We included flags to keep SNPs that are present in a single population (-p 1), required a minimum of 5 reads to call a stack at a locus (-m 5), kept only SNPs (--remove-indels), and export the variant calls in vcf format (--vcf). We then filtered the variant file using Vcftools v0.1.15 (Danecek et al., 2011) for sites with more than 60% completeness (--max_missing .6), calculated the percent missing genotypes for each individual (--missing-indv), and the transition-transversion profiles (--TsTv-summary) for both skin and muscle libraries separately, and then calculated the genome-wide Weir and Cockerham mean Fst between the skin- and muscle-derived libraries (--weir-fst-pop).

### Identification of sex chromosome-specific markers

We used the coverage from each individual to identify sex-linked genomic contigs as described previously (Brekke et al., 2018, 2019) by first calculating each individuals’ sequencing effort as the sum of aligned reads for that individual. We standardised the contig-level counts by dividing by the sequencing effort of each individual and multiplying by 1,000,000. We compared the mean of the coverage of females and the mean coverage of males for each contig. W contigs should be present in females but not males and fulfil the inequality:

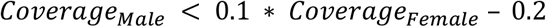

Unknown contigs are the not W-linked and have overall standardized coverage of less than 5:

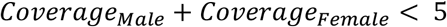

Z-linked contigs are not unknown and have twice the coverage in males as females and fulfil the inequality:

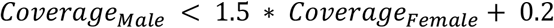

All remaining contigs are annotated as autosomal. These specific cut-offs were chosen based on the natural breakpoints in the plot.

### Chromosomal distribution of SNPs

In late 2019, the DNA Zoo project (https://www.dnazoo.org/), released a chromosome-scale assembly of the corn snake genome, comprising 119,289 contigs (contig N50 37.9kb) in 33,440 scaffolds (scaffold N50 147Mb), assigned to the 18 pairs of chromosomes (8 macro and 10 micro), generated using their 3D *de novo* assembly methodology (Dudchenko et al., 2017). This assembly is based on the initial assembly of Ullate-Agote et al. (Ullate-Agote et al., 2020, 2014). We determined the chromosomal distribution of our SNPs by re-running the stacks pipeline as outlined above on this new chromosome-scale assembly.

## Results

DNA was successfully extracted from all skin and muscle samples, with concentrations varying between 0.37-12ng/µl (mean 6.15ng/µl). In all cases, only a small proportion of a shed skin (typically less than 200mg) was needed to obtain sufficient DNA for GBS. On average there were 2,818,124 ± 1,154,007 reads sequenced per individual. GBS libraries extracted from muscle tissue had more sequenced reads than libraries from skin tissue (Fig. 1a, muscle mean: 3,798,510 reads, skin mean: 2,538,013 reads, Welch two sample T-test, t=4.4064, df=26.183, *P*=0.0001589).

**Figure 1.**
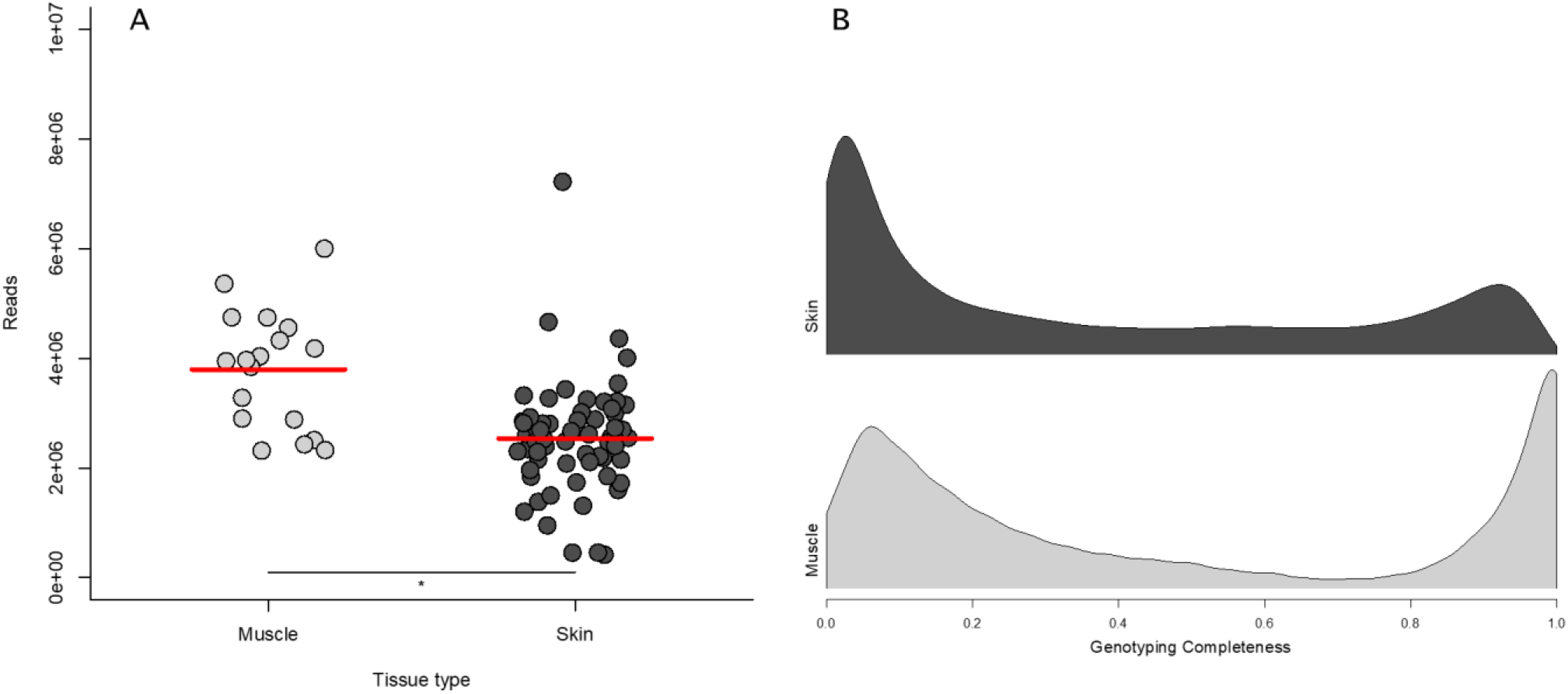
**(a)** Muscle-derived libraries have more reads than skin-derived libraries despite being pooled in equimolar ratios during library construction (Welch two sample T-test, t=4.4064, df=26.183, *P*=0.0001589). This pattern is likely caused by lower quality DNA from dried shed skins than snap-frozen muscle tissue. Each sample is plotted according to its tissue of origin and the red lines denote the mean. **(b)** Density distributions of the proportion of libraries for which each of the 237,466 raw SNPs are genotyped show that fewer sites are genotyped in skin-derived libraries. SNPs at 0 are genotyped in no library, while SNPs at 1 are genotyped in all libraries. In both skin- and muscle-derived libraries there are many SNPs that are only genotyped in a few samples (peaks near 0), but muscle-derived libraries have far more SNPs that are genotyped in many samples (peaks near 1) (T-test, t = 119.47, df = 566235, *P* = 2.2e-16). All further analyses are done on the 101,618 SNPs at completeness of 60% and above.

We identified 237,466 total raw SNPs. After filtering for completeness we found 101,618 SNPs: 97,724 SNPs at >60% completeness in the muscle samples and 70,685 with >60% completeness in the skin samples, of which 66,791 were found in both muscle and skin. Even after filtering for overall missing data, the genotyping rate was highly variable across samples with an average of 16.7% ± 12.7% missing in any given sample, and muscle-derived libraries had fewer missing genotypes than skin-derived libraries (Figure 1b, muscle mean: 11.3% missing, skin mean: 18.2% missing, Welch two sample T-test, t = −3.267, df = 74.723, *P* = 0.001643). Furthermore, there is a strong relationship between the amount of sequencing and the genotyping rate, especially at read counts lower than 1,000,000 (Figure 2).

**Figure 2.**
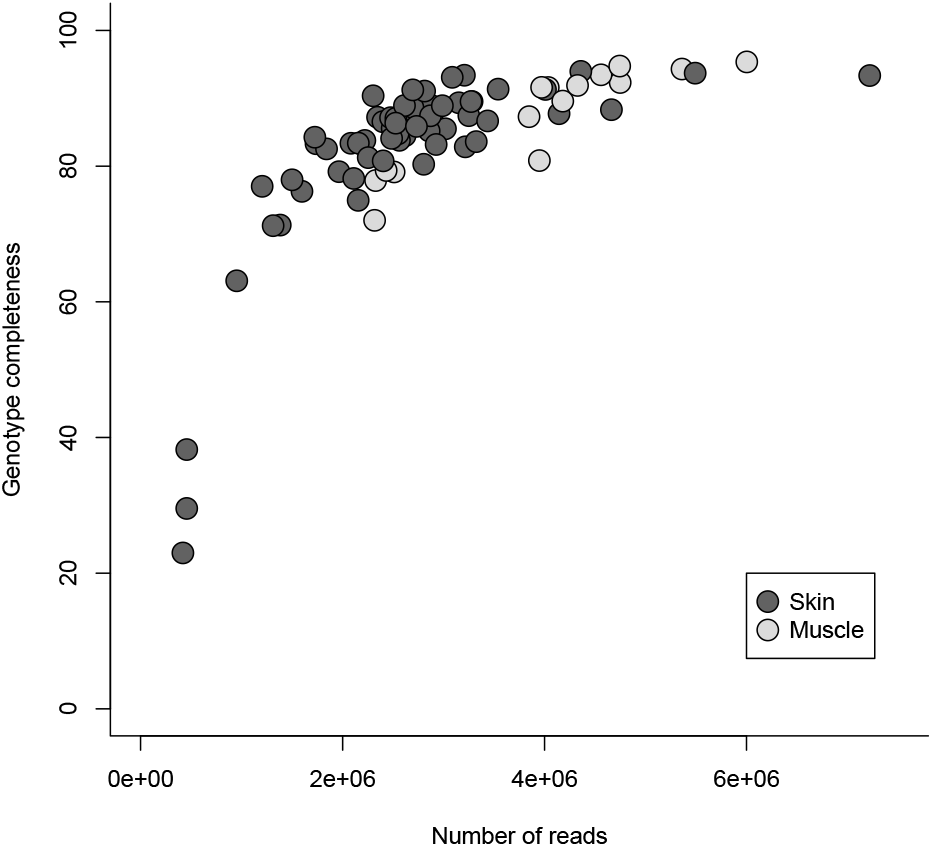
There is a strong relationship between the number of reads sequenced per sample and the number of missing genotypes per individual such that with deeper sequencing more variants can be called. There is a sharp cutoff at around 1,000,000 reads under which the genotype completeness drops sharply, and a consistently high genotyping rate does not occur until above 2,000,000 reads per sample. This figure shows the subset of SNPs at the >60% completeness cutoff for each library type.

Low read counts and much missing data may suggest that the DNA has been damaged prior to library construction. To test for DNA damage we analysed the distribution of transitions and transversions and found significant differences between the skin- and muscle-derived libraries (Figure 3, Chi square test: *X*^2^ = 15.843, df = 5, *P* = 0.007306). While the differences are significant, the effect sizes are slight. The transition to transversion ratio for muscle-derived libraries is 2.652 and for skin-derived libraries it is 2.714 and within each class the two library types differ by only tenths of a percentage: AC in skin is 6.92% and in muscle is 7.07%, AT is 6.57% in skin versus 6.41% in muscle, CG is 6.87% versus 6.92%, GT is 6.86% versus 6.66%, AG is 36.37% versus 36.76%, and CT is 36.24% versus 36.31%. In addition, the Fst between the skin and muscle samples is quite low (Fst = 0.0686) suggesting that there is little genome-wide differentiation between skin and muscle samples.

**Figure 3.**
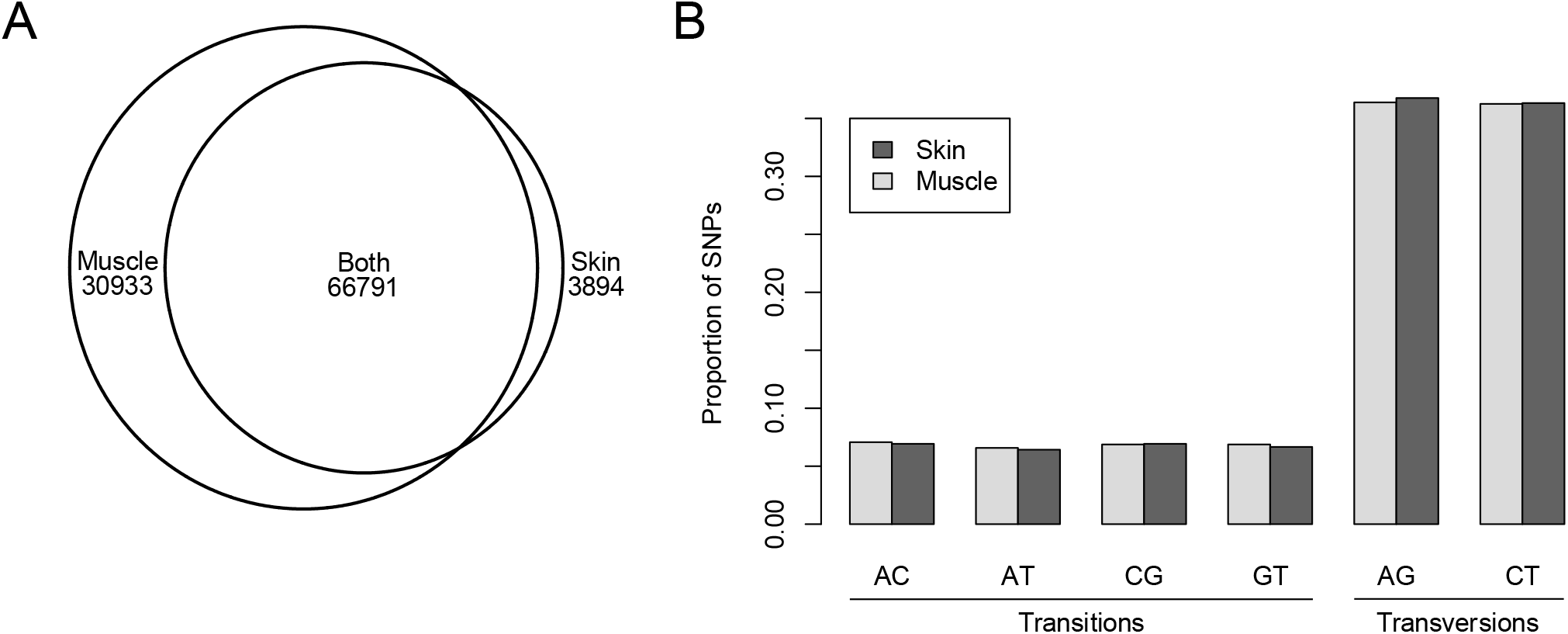
There are slight differences in SNP profiles between skin and muscle libraries. **(A)** After filtering each set of libraries independently for 60% completeness, most SNPs were found in both muscle and skin. **(B)** There are significant differences in the proportions of all SNP types (transitions: AC, AT, CG, and GT, and transversions: AG and CT) between skin and muscle (Chi square test: *X*^2^ = 15.843, df = 5, *P* = 0.007306) but the effect size is vanishingly small. This suggests that DNA damage occurs in the dried skin, but that identifying and accounting for specific errors will be difficult and provide little benefit to the experimental outcome.

We used coverage to identify the sex-linked scaffolds in the 2014 version of the corn snake genome (Ullate-Agote et al., 2014) and were able to reliably annotate approximately 30% of the genome (Figure 4). 58,935 scaffolds (349,080,366 bases) are autosomal, 4178 scaffolds (19,511,706 bases) are Z-linked, and 1275 scaffolds (2,357,273 bases) are W-linked, the remaining 819,528 scaffolds (1,033,270,996 bases) had too little coverage to assign (Supplemental file 1).

**Figure 4.**
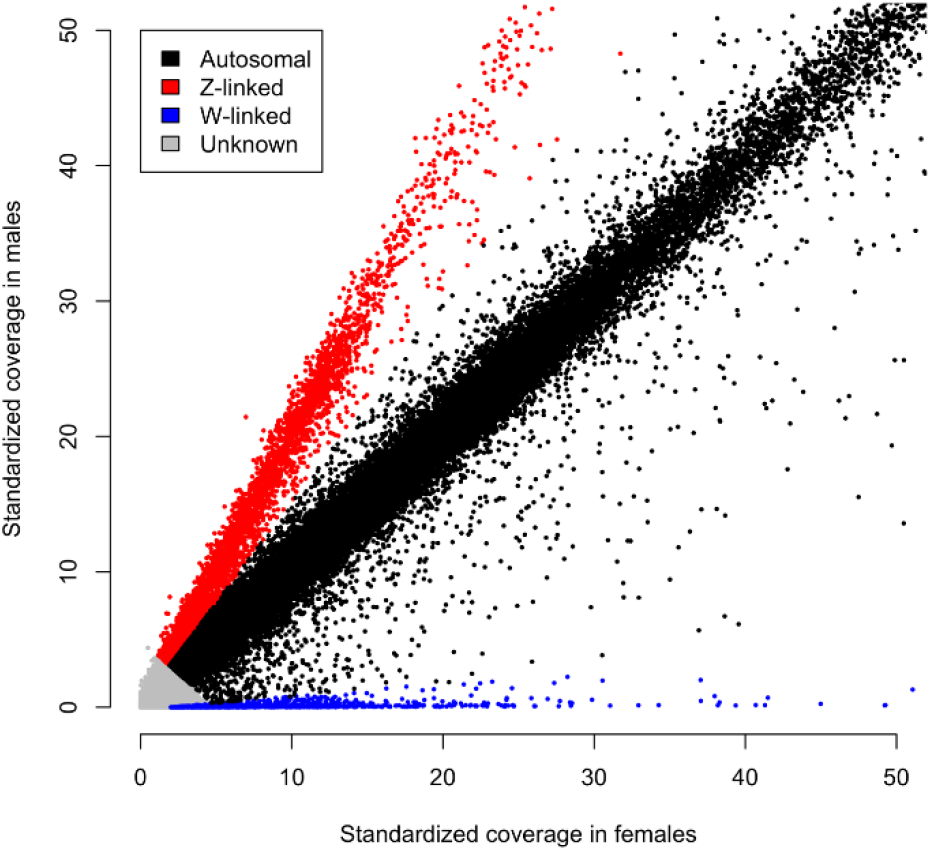
Relative coverage in males and females discriminates sex-linked from autosomal scaffolds. Points on the 1:1 line have equal sequencing coverage in males and females implying autosomal linkage. Z-linked scaffolds (red points) and W-linked scaffolds (blue points) are also readily apparent. Scaffolds with too little coverage to reliably discriminate relative coverage in males and females are shown in grey

When we aligned the 97,638 SNPs with greater than 60% completeness to the recent chromosome-scale corn snake genome assembly (Figure 5), we found that the average SNP density across all chromosomes was 68.13 SNPs/Mb (69.93 SNPs/Mb across all autosomes, and 37.44 SNPs/Mb for the Z), but that there were significantly more SNPs on microchromosomes (mean density 78.39 SNPs/Mb) than macrochromosomes (mean density 55.30 SNPs/Mb (T-test, t = 4.3533, df = 16, *P* =0.000246), Figure 6 and Supplemental Table S2).

**Figure 5.**
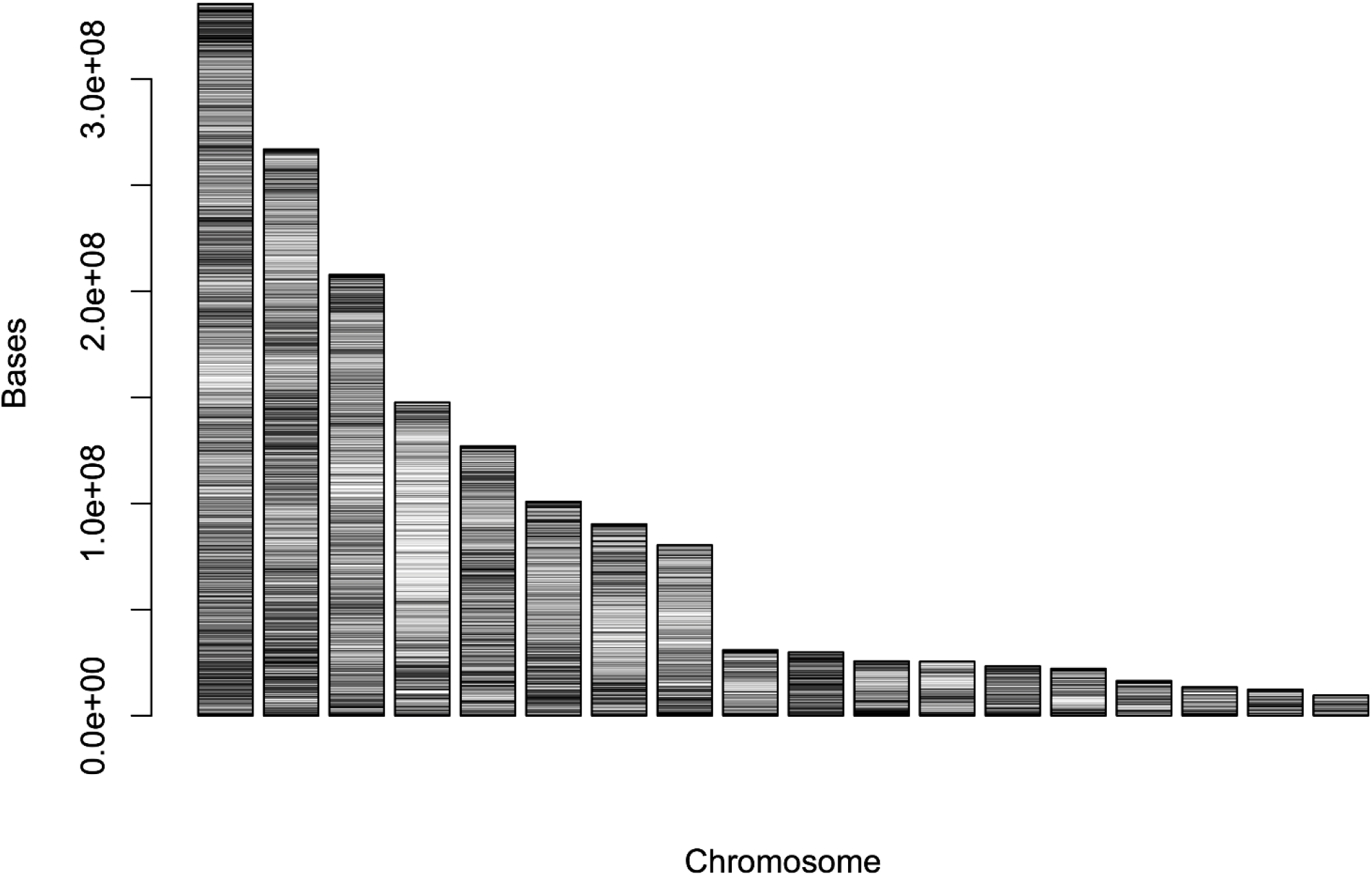
Distribution of SNPs in the corn snake genome. The corn snake has 8 macrochromosomes with an average SNP density of 55.3/Mb, and 10 microchromosomes with an average SNP density of 78.4/Mb. Assignment of chromosome 4 as the Z is based on the localisation of chromosome-specific genes (Matsubara et al., 2006).

**Figure 6.**
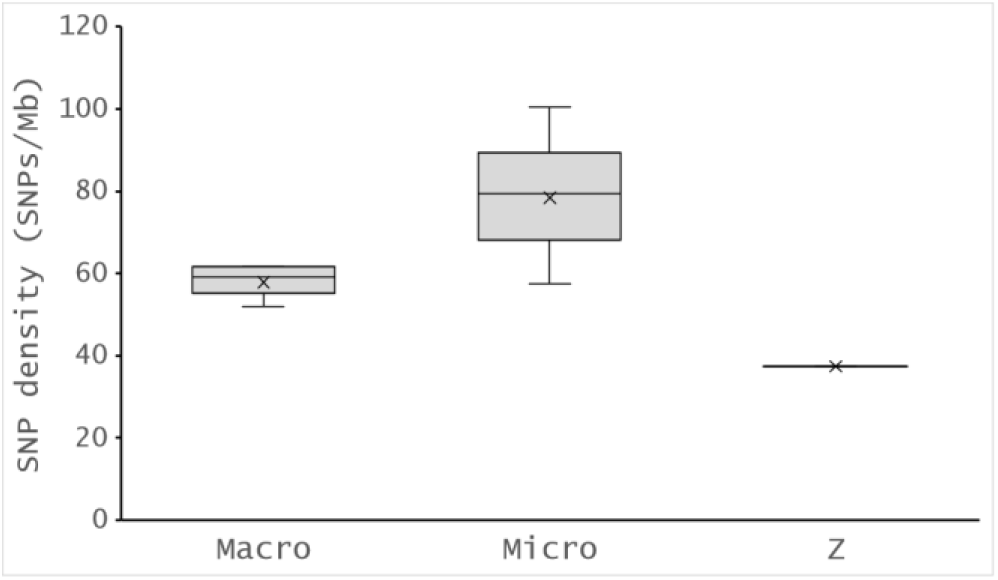
Microchromosomes have significantly more SNPs assigned to them than non-Z macrochromosomes (mean 78.39 SNPs/Mb vs 57.85). The Z chromosome has the lowest SNP density of any corn snake chromosome in this study (37.44 SNPs/Mb)

SNP density tends to scale with chromosome length, with the exception of the Z chromosome, which has fewer SNPs than would be expected given its size (37.44 SNPs/Mb vs the non-Z macrochromosome average 57.85 SNPs/Mb and all-autosome average 69.93 SNPs/Mb), and the (micro)chromosome 10, which has the highest SNP density in the corn snake genome (100.51 SNPs/Mb) (Figure 7).

**Figure 7.**
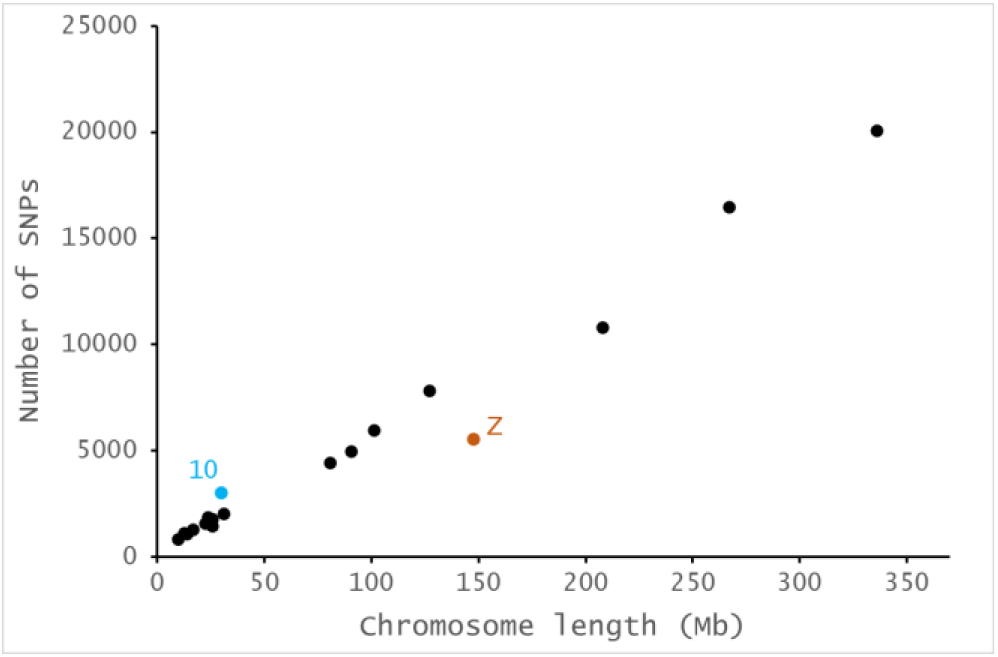
SNP density scales with chromosome length, with the exception of the Z chromosome (orange), which has fewer than expected, and microchromosome 10 (blue), which has the highest SNP density in the corn snake genome (100.51 SNPs/Mb, see Supplemental Table S2).

## Discussion

DNA extracted from shed skins is suitable for use with reduced-representation sequencing approaches such as GBS with some considerations. We compared GBS libraries built from skin-extracted DNA with libraries from muscle-extracted DNA. The muscle samples were immediately snap-frozen and are thus a source of high-quality DNA while the skins were collected by pet snake owners across the country, dried, and shipped at ambient temperature through the post and thus subject to a variety of mechanisms of DNA degradation. In this way they more accurately reflect the type of material that may be collected from captive specimens, or what may be obtained via citizen science projects such as the ‘Reptile Slough Genebank Project’. We found significant differences between the muscle- and skin-derived DNA that likely stem from DNA damage in the original samples. These differences include the number of reads sequenced (Figure 1a) and SNPs identified (Figure 3). However, dealing with the differences between the two library types is surmountable with some forethought toward experimental design.

The number of SNPs identified in a GBS experiment depends strongly on the sequencing depth (Figure 2). Thus, sequencing skin samples more deeply than samples from fresh tissue will help assure that a sufficient number of real SNPs can be identified. Estimating the necessary coverage in a GBS experiment is difficult as it depends on the frequency of cut-sites in the genome (often unknown in non-model species), the heterozygosity present in the sample population (often unknown until at least after the first round of sequencing), the specific cut-offs used to filter variant sites, and the type of experiment (population genetics may require different number of SNPs than parentage analyses etc). For this experiment the SNP discovery curve plateaued at around 3,000,000 reads per individual (Figure 2), and many of the skin-derived libraries had far fewer genotyped SNPs due to low read coverage (Figure 1b and 2). As such, we suggest that future researchers plan on sequencing skin and other possibly damaged samples more deeply in order to identify a robust set of variants.

Some problems, especially those relating to DNA damage, cannot be dealt with simply by sequencing more deeply. For instance, if Cytosine deamination into Uracil is a common issue in the sample (as is often the case in ancient DNA studies (Hofreiter et al., 2001), more sequencing will not remove those errors and a more sophisticated approach is needed. The signal of errors in the skin libraries is very slight suggesting that identifying and dealing with these errors would be difficult. Fortunately, such labour-intensive error cleaning will likely provide little benefit for three reasons. First, the vast majority of the SNPs identified in the skin samples are also found in the muscle samples (Figure 3a) and this is true despite the excess of skin-derived libraries (61 vs 18 muscle-derived) in which to discover SNPs. Even if the 3,894 skin-specific SNPs are not biologically real, they are so few (only 3.8% of all SNPs) that that identifying them in the absence of other high-quality libraries will be exceedingly difficult and more importantly, removing them will have little effect on the overarching results. Secondly, while the skin and muscle may have slightly different SNP profiles (Figure 3b), the effect is so slight that filtering options such as removing all C/T sites or ignoring all transversions are not merited. Finally, if there was a systematic damage pressure, it should alter the allele frequencies in the damaged samples. But the Fst between the skin and muscle samples is very low (genome wide mean Fst is 0.0686). While this metric can only be calculated for the 66,791 shared SNPs, it shows that there are not striking allele frequency differences between skin- and muscle-derived libraries which is evidence that the degradation is not a major concern. Such a low Fst result is additionally convincing given that the individuals from whom muscle was used are all close relatives in our colony while the snakes whose skins were used originate from breeders across the UK. This confounding population structure should artificially increase our estimate of Fst, and the Fst due solely to DNA damage in the skin is likely much smaller. In sum, while DNA damage is apparent in our skin samples, the effects are slight and not likely to impact any biological results.

The SNPs we identified are not distributed evenly across the corn snake genome, although SNP density does generally correlate with chromosome length (Figure 7). Microchromosmes have significantly higher SNP density than macrochromosomes (Figure 6), which should perhaps not be too surprisingly given the well-known differences in base composition between macro and microchromosomes (Srikulnath et al., 2021). Indeed, in the corn snake, macrochromosomes have an average GC% of 39.69% and microchromosomes 44.79% (Supplemental Table S2). The PstI enzyme we used in preparation of our GBS libraries has a GC-rich cut site (CTGCA^G) and so is likely to cut more often on microchromosomes. Future researchers may therefore wish to consider the GC-richness of microchromosomes when choosing restriction enzymes for reptile GBS experiments.

Leveraging the amateur herpetological community and sourcing skins from the public has the dual benefit of engaging citizen scientists and the possibility of rapidly collecting extremely large sample sizes. Pet snakes in general, and corn snakes in particular, have a variety of colour and pattern morphs which may prove incredibly powerful for understanding the genetic basis for colouration (Ullate-Agote et al., 2020, 2014). We have shown that a simple crowd-sourced collection technique (mailing shed snake skins) can provide samples containing DNA of sufficient quality for reduced representation sequencing. This finding opens up the possibility of doing association studies on patterning and colouration from a wealth of samples in a long-lived and low-fecundity species, although more research is required into the impact of time since shedding, and UV exposure (especially for field-collected sheds) on DNA quality.

## Acknowledgements

We wish to thank Adam Clarke and numerous anonymous donors for both financial support and donation of shed skins. We also wish to thank the School of Natural Science technical team, and especially Rhys Morgan, for assistance with animal care, and Adam Hargreaves for useful discussions.

## Data Accessibility

The datasets generated during and/or analysed during the current study are available in the European Nucleotide Archive (ENA) under project PRJEB32869.

## Author contributions

JFM devised the study. LS and MJH performed experiments, and TDB and JFM analysed the data and wrote the paper.

## Supplementary tables

**Supplementary Table S1.**
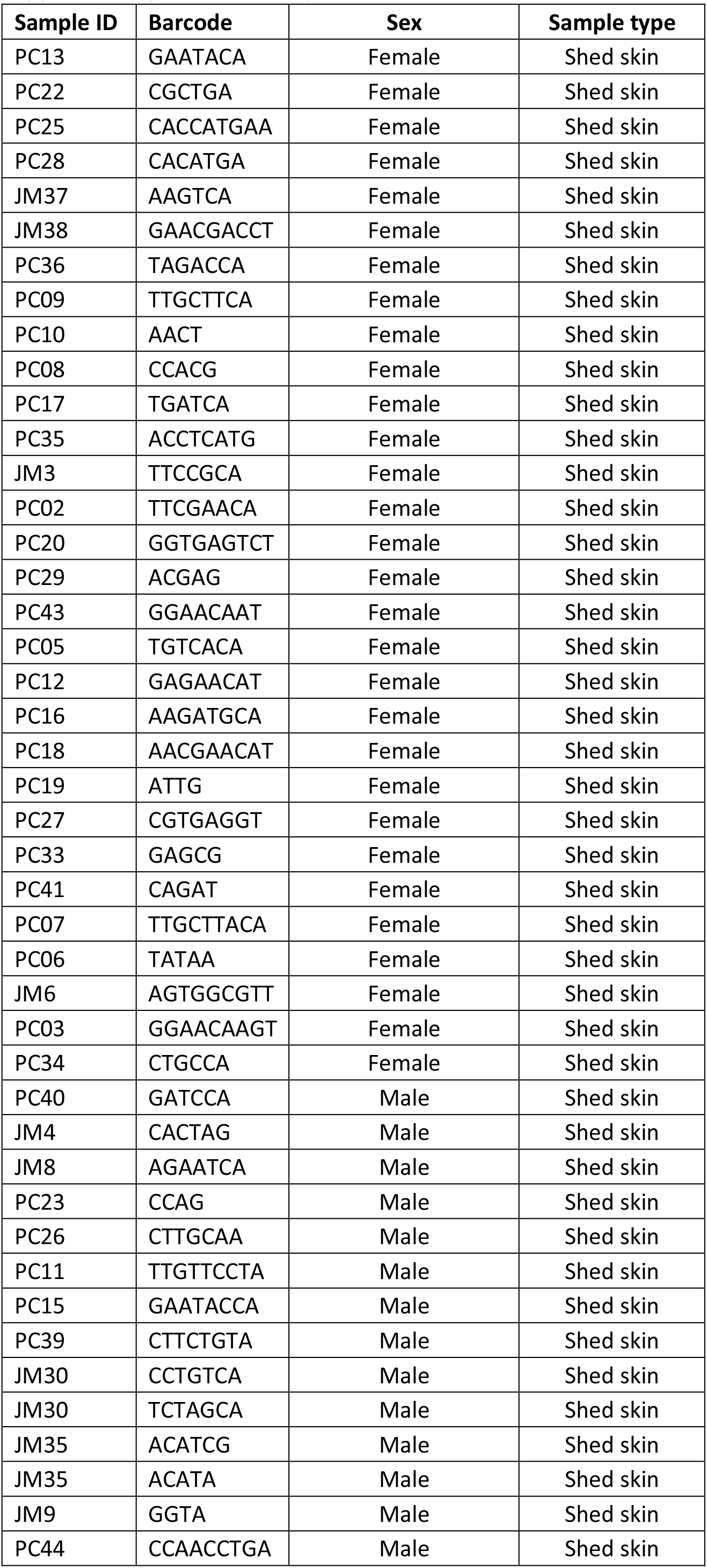
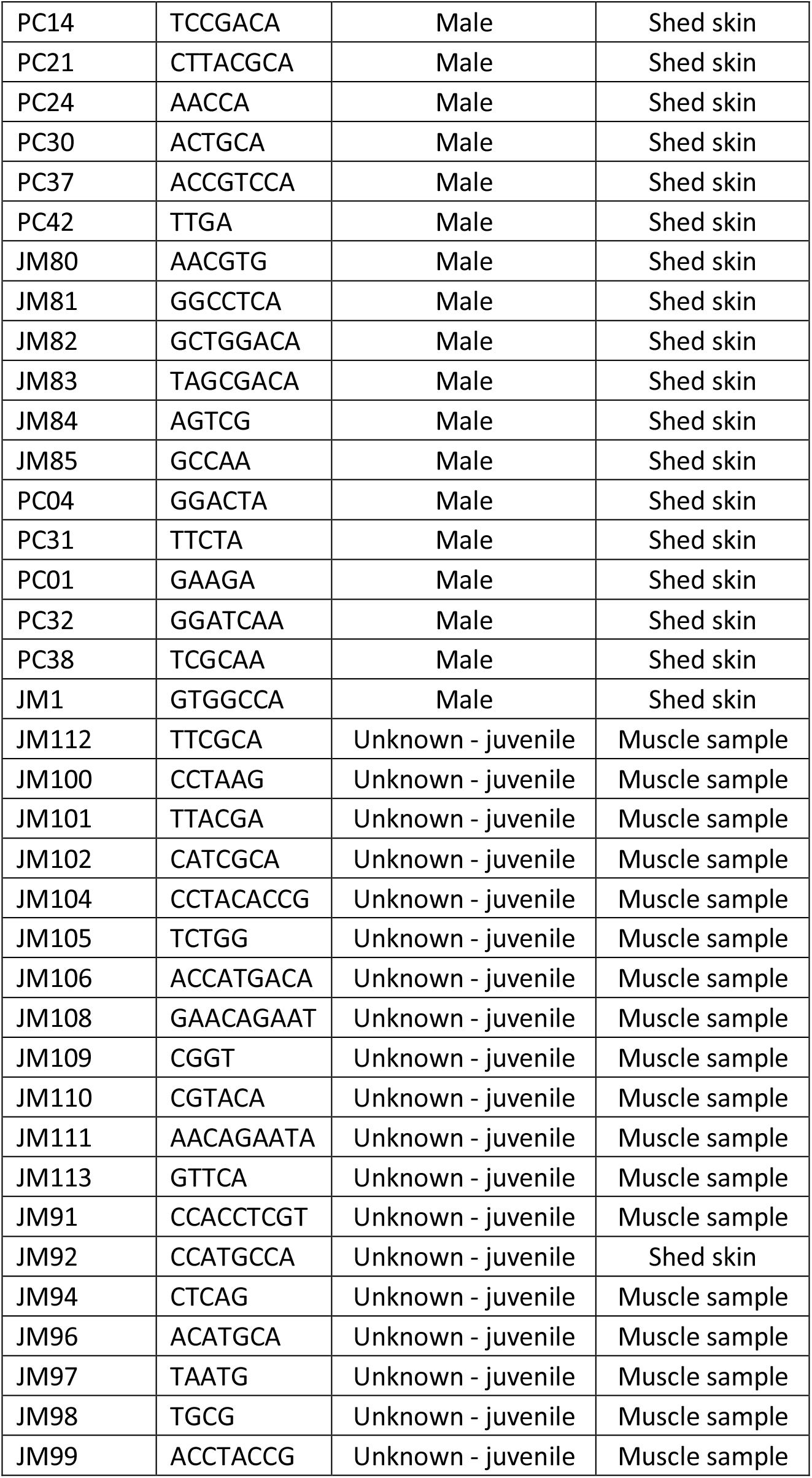
Sample details.

**Supplementary Table S2.** Distribution of SNPs in the corn snake genome.

|  | Chromosome | Number of SNPs | Chromosome length (Mb) | SNP density (SNPs per Mb) | GC% |
| --- | --- | --- | --- | --- | --- |
| Macro | 1 | 20070 | 335.6 | 59.80 | 39.21 |
|  | 2 | 16490 | 267.0 | 61.77 | 39.64 |
|  | 3 | 10794 | 207.8 | 51.94 | 39.12 |
|  | 4 (Z) | 5526 | 147.6 | 37.44 | 40.02 |
|  | 5 | 7831 | 127.0 | 61.65 | 39.53 |
|  | 6 | 5978 | 100.9 | 59.23 | 39.58 |
|  | 7 | 4977 | 90.3 | 55.11 | 40.40 |
|  | 8 | 4454 | 80.3 | 55.45 | 40.00 |
| Micro | 9 | 2022 | 30.9 | 65.34 | 42.97 |
|  | 10 | 3000 | 29.8 | 100.51 | 44.08 |
|  | 11 | 1772 | 25.7 | 68.94 | 42.95 |
|  | 12 | 1464 | 25.5 | 57.45 | 43.08 |
|  | 13 | 1862 | 23.3 | 80.00 | 43.72 |
|  | 14 | 1570 | 22.2 | 70.78 | 44.82 |
|  | 15 | 1295 | 16.4 | 78.76 | 45.93 |
|  | 16 | 1093 | 13.4 | 81.30 | 46.23 |
|  | 17 | 1138 | 12.3 | 92.29 | 47.00 |
|  | 18 | 847 | 9.6 | 88.55 | 47.09 |

Summary statistics:
|  | SNP density |  | Mean GC% |
| --- | --- | --- | --- |
|  | Mean | Median |  |
| All chromosomes | 68.13 | 63.55 | 42.52 |
| All macrochromosomes | 55.30 | 57.34 | 39.69 |
| Macrochromosomes (no Z) | 57.85 | 59.23 | 39.64 |
| Microchromosomes | 78.39 | 79.38 | 44.79 |
| All autosomes | 69.93 | 65.34 | 42.67 |

